# Fast and light efficient remote focusing for volumetric voltage imaging

**DOI:** 10.1101/2023.11.28.568783

**Authors:** Urs L. Böhm, Benjamin Judkewitz

**Affiliations:** Einstein Center for Neurosciences, Charité – Universitätsmedizin Berlin, Germany

## Abstract

Voltage imaging holds great potential for biomedical research by enabling noninvasive recording of the electrical activity of excitable cells such as neurons or cardiomyocytes. Camera-based detection can record from hundreds of cells in parallel, but imaging entire volumes is limited by the need to focus through the sample at high speeds. Remote focusing techniques can remedy this drawback, but have so far been either too slow or light inefficient. Here, we introduce FLIPR, a new approach for remote focusing that doubles the light efficiency and enables high-speed volumetric voltage imaging at 500 volumes/s. We show the potential of our approach by combining it with lightsheet imaging in the zebrafish spinal cord to record from >100 spontaneously active neurons in parallel.

## 1. Introduction

Functional imaging with genetically encoded indicators of neural activity can reveal detailed insights into the inner workings of the nervous system. Light sheet imaging in combination with genetically encoded voltage indicators has shown great potential to record the membrane potential of tens of neurons in parallel [1,2], but due to the high acquisition speeds necessary for voltage sensors (500-1000 Hz) imaging is usually limited to a single focal plane. One major reason for this limitation is the requirement to focus through the sample at sufficient speed. Since most neural tissue exists in 3D, it is desirable to have microscopy methods that can image from a volume instead of a single focal plane.

A common technique for focusing through a sample in light sheet microscopy is to move the imaging objective axially with a piezo-electric actuator, but due to the inertia of relatively heavy objectives this is limited to a few tens of Hz at best. An alternative approach is to use remote focusing, which places the focusing element away from the primary objective and the sample. Remote focusing can be achieved in two ways: either by introducing defocus in a Fourier plane of the imaging system by using a tunable focusing element [3–7] or by refocusing a remote image plane (remote focusing) [8–10]. The latter method has the advantage that it can be used to achieve fast aberration free imaging over a relatively wide z-range away from the nominal focal plane of the primary objective [8]. However, by using a small movable mirror in the remote image plane (Fig 1a, left), this method has the disadvantage of losing >50% of light due to the necessity of a quarter-wave plate and polarizing beamsplitter (PBS) to separate incoming and refocused light. This is especially harmful in ultra-high-speed recordings such as voltage imaging, where integration times are in the sub-millisecond range and the signal-to-noise ratio (SNR) is severely limited by the amount of photons that can be collected in each frame. Remote focusing has therefore mainly been used to achieve fast axial scanning in two-photon microscopy [9,10] and is not widely adopted as a method on the emission light path to refocus the detected image [11–13].

**Fig. 1.**
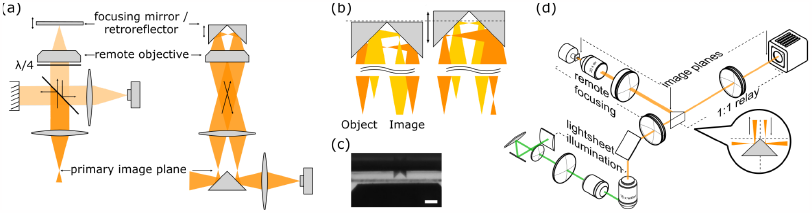
Principle of the approach: (a) Conceptual layout of our proposed design. Current polarizing beamsplitter based designs (left) lose half of the light intensity because unpolarized fluorescence is split into a beam dump and the remote focusing objective. Our design (right) uses half the available FOV for the incoming light and a retroreflector in the image plane of the remote objective to fold the image to the other side of the FOV. A knife edge mirror in the primary image plane is then able to direct the refocused light onto the camera. (b) Light path at the level of the object, retroreflector and image of two point sources located at different z-depths. Moving the retroreflector along the optical axis brings either of them in focus. (c) Picture of the microscopic retroreflector together with the coverslip and parts of the remote objective. Scale bar = 500 μm. (d) Layout of the entire microscope for volumetric voltage imaging. A light sheet is generated with a cylindrical lens and projected onto the sample via a scan and tube lens and illumination objective. Its z-position is controlled by a galvo mirror. Fluorescence is collected via the imaging objective and imaged onto the knife edge mirror in the primary image plane and directed into the remote focusing path. After refocusing the image is relayed with a 1:1 relay onto the camera.

Here, we propose a new remote focusing design that doubles the light efficiency, called FLIPR (flipped image remote focusing). FLIPR maintains the advantage of high-speed remote focusing, but does not rely on polarization optics to separate the incoming from the refocused image. Instead of placing a mirror in the remote image plane, we use a microscopic retroreflector to flip and fold the image back into the remote focusing arm and spatially separate incoming and outgoing light (Fig 1a-c). This design achieves volumetric imaging at up to 500 Hz for a z-depth of 150 μm. We show its potential for enabling new high-speed volumetric imaging applications by recording the membrane potential of over a hundred neurons in the spinal cord of larval zebrafish in parallel.

## 2. Results

To separate the refocused light from the incoming light in a remote focusing system, previous designs typically used a polarizing beamsplitter with a quarter-wave plate, which is light inefficient due to the fact that fluorescence is unpolarized (Fig. 1a). FLIPR circumvents the need for polarization optics by making use of a miniature retroreflector. In addition to shifting the focal plane, the retroreflector flips half of the image from one side of the field of view (FOV) of the remote objective to the other side. The refocused image can be picked off and relayed to the camera with a knife edge mirror in the conjugate image plane (Fig. 1b).

Based on this approach, we built a remote focusing system using a 16x 0.8 numerical aperture (NA) water dipping primary objective and a 20x 0.75 NA air remote objective. By using a non-standard focal length of 124 mm for the remote system tube lens, we achieved near-unit angular magnification of M = n_1_/n_2_ ≈ fO_1_/fO_2_ = 1.29 (with n_1/2_ being the respective refractive index of the immersion medium of the primary and remote objective) [8]. To achieve the proposed flipping of the image at the remote focusing plane, we custom built a microscopic retroreflector from two 0.5 mm aluminum coated right-angle prisms. To determine the performance of our setup, we measured the point spread function (PSF) at multiple positions along the z-axis and quantified lateral resolution as well as maximal intensity. Our setup achieves a maximal lateral resolution of 0.54 μm (theoretical limit: 0.34 μm) and maintains 80% of its maximal intensity over a z-range of approximately 100 μm (Fig. 2ab).

**Fig. 2.**
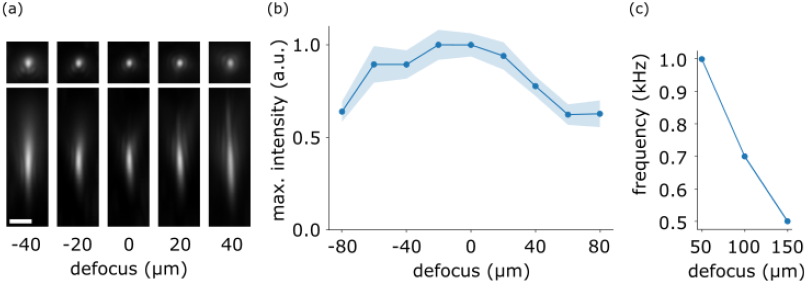
Optical characterisation. (a) Measured point spread function from 0.1 μm diameter fluorescent beads over a 80 μm defocus range. Scale bar: 2 μm. (b) Average maximal intensity from each PSF normalized to the signal at 0 defocus. Intensity stays >0.8 over a range of approximately 100 μm. Shaded area designates standard error of the mean (s.e.m.), n = > 39-103 beads for each data point. (c) Maximum drive frequency limited by the voice coil motor as a function of defocus.

Using a linear voice coil motor, we were able to move the retroreflector along the z-axis in the sample plane of the remote objective with up to 500 Hz over a range of 150 μm (Fig. 2c). This allowed us to access a volume of 388 μm in x-direction (limited by the size of the retroreflector) and 150 μm along the z-axis while the y-axis is limited by the speed of the camera essentially trading off FOV along the y-axis and number of planes in the volume. Due to inertial forces during high speed motion of the voice coil motor, the retroreflector does not follow a strictly linear trajectory along the optical axis, but rather moves along an ellipsoid (Fig. S1ab). This results in a reproducible shift of ≤ 6.5 μm between the up- and the down-stroke along the y-axis of the image, which can be compensated digitally (perpendicular to the axis along which the image is flipped by the retroreflector, Fig. S1c). Due to the continuously moving refocusing system and the rolling shutter of the camera, optical planes are also tilted in the volume relative to z-axis and in opposite directions during the down- and up-focus part of the volume (Fig. S2a and Fig. 3a). Both y-movement and tilt have to be taken into account for data analysis (Fig. S2b).

**Fig. 3.**
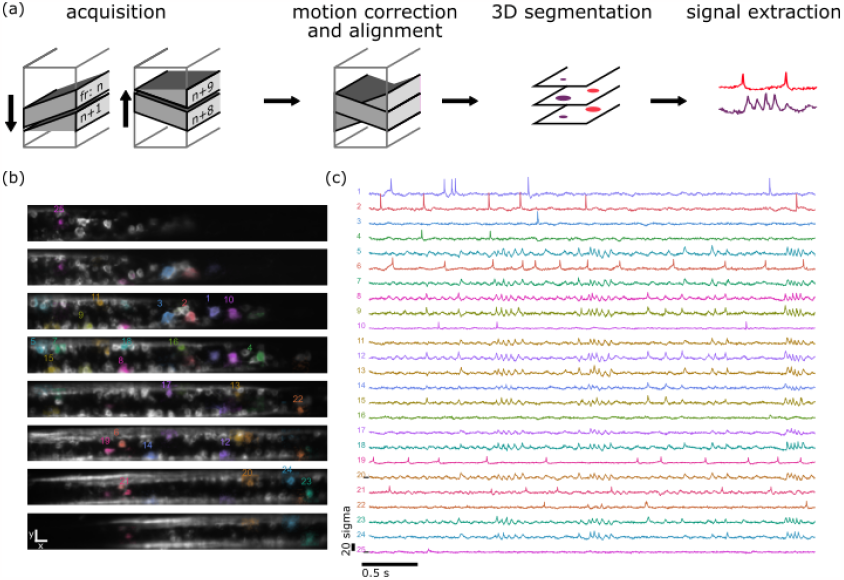
High-speed volumetric voltage imaging: (a) Schematic of the pipeline for volumetric voltage imaging data acquisition and signal extraction. 8 frames are acquired during the downward and another 8 during the upward stroke of the refocusing cycle. Average frames at each position are then motion corrected and aligned to generate one single volume from which individual cells are segmented in 3D. The obtained 3D ROIs are then used as the basis for the extraction of the fluorescent time traces (see methods for details). (b) Average fluorescence of 8 planes spanning the entire volume of 50 μm of spinal cord tissue of a 4 dpf zebrafish larva. animals are expressing UAS:Voltron2-ST under the control of HuC:Gal4 and are stained with JF-526 dye. Footprints of select neurons shown in (c). Most footprints are visible over several z-sections but numbered only once. scale bar 20 μm. (c) Fluorescent traces corresponding to the spatial footprints shown in B (z-scored). Several neurons show clear single spikes while others show distinct oscillations likely corresponding to fictive swimming activity.

To showcase the performance of FLIPR for volumetric voltage imaging, we imaged the activity of spinal cord neurons in zebrafish larvae expressing the voltage indicator Voltron2 (*Tg(HuC:Gal4; UAS:Voltron2-ST)*) [14]. We recorded 20 s of spontaneous activity in a volume spanning approximately 4 spinal segments (x = 390 μm, y = 46 um, z = 50 μm) divided into 2x8 planes (up-stroke and down-stroke) at 500 volumes/s. The data was then aligned and motion corrected and cell bodies were segmented in 3D to identify >200 cell bodies. After signal extraction and manual curation (see methods) the dataset contained meaningful signals from >100 neurons (Fig. 3b,c and Fig. S3). Among the recorded neurons, some showed clear rhythmic membrane oscillations expected during fictive locomotor behavior, whereas others showed seemingly independent single spike activity (See also Video S1).

## 3. Discussion

Random access scanning [15] and multiplane confocal scanning [16] have recently attempted to expand voltage imaging to 3D volumes but remain limited by either the number of cells that can be recorded in parallel or the accessible volume owing to the limitations of point scanning approaches. Here we developed FLIPR, a technique that combines a light efficient optical design for remote focusing with the advantages of high speed camera based readouts to potentially measure hundreds of neurons from large volumes. This is achieved by sacrificing half the FOV to obtain the light efficiency necessary for voltage imaging (see [17] for an alternative approach). Since current camera technology cannot image the entire FOV at the kHz rates necessary for volumetric voltage imaging, decreasing the FOV should not be considered a drawback. It should be noted that the method described here is not limited to light sheet microscopy; rapid and light efficient refocusing could also be used in widefield microscopy e.g. to provide volumetric data during voltage imaging from mouse cortex or any other application where speed and light efficiency cannot be traded off for each other.

Depending on the nature of the sample, the volume from which we can record at a given rate is limited either by the camera speed or the maximal force output of the voice coil motor. For sparsely labeled samples, the refocusing can happen over a larger z-range within a single camera frame without projecting neurons on top of each other and thereby decreasing signal-to-noise. In such a situation the accessible volume is limited by the maximal range that the voice coil motor can scan over at a given frequency (which is limited by the maximal acceleration and thus the force output of the device). For very dense samples, where relatively fine grained sampling along the z axis is necessary to avoid projecting cells on top of each other, the main limitation becomes the frame rate of the camera. This consideration also shows that the total number of recorded neurons will be approximately the same in both scenarios as it is ultimately only a function of camera data rate.

Due to the 90° opening angle of the retroreflector and the unit angular magnification, our design limits the effective NA of the primary objective along one axis to NA < 0.7n (with n = refractive index of the immersion medium of the primary objective). This was not a limitation in the setup used here. Lastly, the FOV along the x-axis is currently limited by the size of the retroreflector. A custom non-square prism that is wider but not higher (to allow enough space to move within the limited working distance of the remote objective) would increase the FOV along the x-axis without compromising the FOV in y and z.

For technical simplicity, the illumination light in our setup stays constant over the entire period of focusing through the volume. To avoid having to align differently tilted and laterally shifted images from the up and down stroke phase of the volume scan, shuttering the light off during either of the phases would simplify data analysis significantly. Analog modulation of laser intensity could also be used to correct for differences in z-speed (and thus integrated illumination intensity) of the lightsheet at the center of the volume versus the turnaround points. This can also be mitigated by using a more powerful voice coil motor to better approach a triangle or sawtooth wave instead of the heavily filtered triangle/sinusoid intermediate used in our current setup. With these improvements, we anticipate that FLIPR can be used to image hundreds of cells at kHz rate.

## 4. Methods

### Microscope setup

Light Sheet illumination path: The lightsheet was generated from a 532 nm laser (CNI MSL-FX-532), passed through a cylindrical lens (Thorlabs LJ1878L1-A, f = 10 mm) and beam reducer (f1 = 100 mm, f2 = 75 mm). Z-scanning was done with a galvo mirror (Cambridge Technology 6215H) before sending the beam through a scan lens (f = 50 mm), tube lens (f = 400 mm), and objective (Olympus XFluor 4X, NA 0.28) into the imaging chamber. This resulted in a nominal Rayleigh length of the lightsheet of 17 μm and 1/e^2^ width of 3.4 μm. To eliminate low amplitude, high-frequency intensity fluctuations from the galvo position, focal length of all lenses was chosen to maximize the angle the galvo has to move to generate a given z-movement of the lightsheet while still maintaining the desired width and Rayleigh length.

Detection and remote focusing path: Fluorescence was detected with a 16x 0.8 NA water immersion objective (Nikon LWD 16x, NA 0.8) and filtered with a longpass filter to reject light from the excitation laser (Chroma ET542LP). The primary tube lens consisted of a 200 mm Plössel lens made from two 400 mm achromats (Thorlabs AC508-400-A). At the primary imaging plane the light was sent into the remote focusing arm using a knife edge mirror. The remote focusing arm consisted of a secondary tube lens made of a 180 mm tube lens (Thorlabs TTL-180) and an additional 400 mm achromat (Thorlabs AC508-400-A) to achieve an effective focal length of 124 mm. As a remote secondary objective we used a 20x 0.7 NA air objective with a coverslip (Nikon Plan Apo 20x, 0.7 NA, WD = 1 mm) to achieve an overall magnification of M = 1.29, close to the ideal M = 1/n = 1.33. The microscopic retroreflector was custom built from two aluminum coated 45 degree prism mirrors (Toweroptical 4531-0020) glued onto an acrylic support structure and mounted on a linear voice coil motor (Rapp Optoelectronic GLP-V1. After refocusing, the image was relayed with two 300 mm lenses (Thorlabs AC508-300-A) in Plössl configuration onto a high speed sCMOS camera (Teledyne Photometrics Kynetix).

To assess the maximal optical performance, the setup was simulated in Zemax to determine the diffraction limited FOV and with code published in [18] to simulate the remote focusing range.

### Data acquisition

The camera was controlled using the manufacturer’s proprietary software while all other waveforms to control the voice coil motor and galvo mirror were generated with custom python code.

The alignment and synchronization between lightsheet and remote focusing system was calibrated by roughly defining the top and bottom position of both lightsheet and focusing system in the volume of interest to get a rough initial alignment. The remote focus was then positioned at 10 equidistant focal planes in that volume and the 31 pictures with varying lightsheet position around that focal plane were acquired. The ideal lightsheet position was then determined by finding the plane with the highest contrast as described here [19] and linear regression was used to find the final relationship between lightsheet and galvo position. During high-speed recordings we noticed a slight phase lag of the voice coil motor relative to the galvo mirror that led to some planes being out of focus. This was corrected by adding a manually determined phase shift. The galvo and voice coil motor were driven with a 500 Hz triangle wave to ensure the maximum time of linear motion through the volume. To avoid too much strain on the voice coil motor, the waveform was low pass filtered at 2000 Hz. The camera was run in ‘frame-overlap trigger’ mode where the exposure time of each frame is determined by the high state of an incoming trigger pulse to ensure that each frame would expose during the same phase of the volume scan.

Data was acquired at 500 Hz volume rate with 16 frames per period (8 frames per sweep through the volume). The camera was therefore triggered at 8 kHz.

### Optical characterization

To acquire point spread functions (PSF), the main camera was replaced by a camera (model) with smaller pixel size to better sample the PSF at the diffraction limit. Stacks of 0.1 nm beads suspended in 1.5% agarose were acquired and a Gaussian was fitted to the x-, y-, and z-projection of each bead. The mean position of the fitted Gaussian was used to center an upsampled version of the bead image before averaging individual PSFs within an 127 × 58 x 12.6 μm^3^ volume. The peak fluorescence reported in Fig. 2b is the average peak value of the Gaussian fit to the x-profile of each bead.

### Larval zebrafish voltage imaging

3 days post fertilization (dpf) *Tg(HuC:Gal4; UAS-Voltron2-ST)* zebrafish larvae were stained for at least 2 h in a solution of 3 μM Janelia Fluor 532 dye + 3% DMSO in fish facility water, rinsed twice and left to wash out remaining dye for at least 1h. 3-4 dpf larvae were paralyzed by immersion in 1 mg/ml α-bungarotoxin for 2-3 minutes and mounted in 1.5% low melting point agarose. To increase overall activity in the nervous system, the imaging chamber contained 20 mM pentylenetetrazole (PTZ). Maximal laser intensity after the lightsheet objective was 15 mW.

### Data processing

Due to a mismatch between the DAQ clock generating the frame trigger and the camera internal clock, we noticed a jitter in the exposure time of each frame that led to small changes in full frame pixel counts as well as changes to baseline pixel values of each line which introduced additional noise. We first detected the timing of these changes and corrected for the relative change in fluorescence due to exposure time differences as well as additive changes in baseline for each pixel row.

Time series for single planes were then motion corrected using the motion correction module included in the Caiman package [20].

To generate a single 3D volume to draw regions of interest, the average fluorescence of each plane was first aligned to its adjacent planes. The rolling shutter nature of the camera results in a slightly tilted plane relative to the optical axis and the tilt is in the opposite direction when moving through the volume from top to bottom as compared to from bottom to top. To combine all planes in one single 3D stack for segmentation, each plane was therefore divided in half and stitched with the other half of a plane most closely matching its actual z-position (see Fig. S2b for an illustration).

The resulting 3D stack was then automatically segmented to generate 3D regions of interest (roi) using cellpose and a custom trained model [21].

To extract fluorescent time traces from the volume, we adapted the method described by Cai et al. [22] to be used with 3D data instead of single plains. Briefly, this method estimates an initial spike train from the average fluorescence of the roi and a background signal from a region around the roi. Multiple iterations then produce a weighted sum of pixel values to only include fluorescence that contributes to the signal from a given cell. All traces and spatial footprints were manually checked to remove obvious duplicates and traces without neural activity.

All custom code for microscope control and data analysis will be made public upon publication.

## Back matter

### Funding

We acknowledge support by the German Research Foundation (DFG, projects EXC-2049-390688087 and 432195732) the European Research Council (ERC2016-StG-714560, ERC2021-CoG-101043615), the Einstein Foundation (EPP-2017-413), and the Alfried Krupp von Bohlen und Halbach Foundation.

## Acknowledgments

We thank C. Berlage for initial discussion of the optical layout, M. Hoffmann, V. Cook, M. Kadobianskyi, J. Veith and C. Berlage for critically reading the manuscript. We thank E. Schreiter for sharing the UAS:Voltron2-ST line and M. Renz, A. Wrana and N. Kroworz for fish care.

Data analysis was performed on the Berlin Institute of Health high-performance compute cluster.

## Disclosures

The authors declare no conflicts of interest.

## Data availability

The raw data underlying the results presented in this paper will be made available upon publication.

## Supplemental document

See Supplement 1 for supporting content.

---

**Figure S1.**
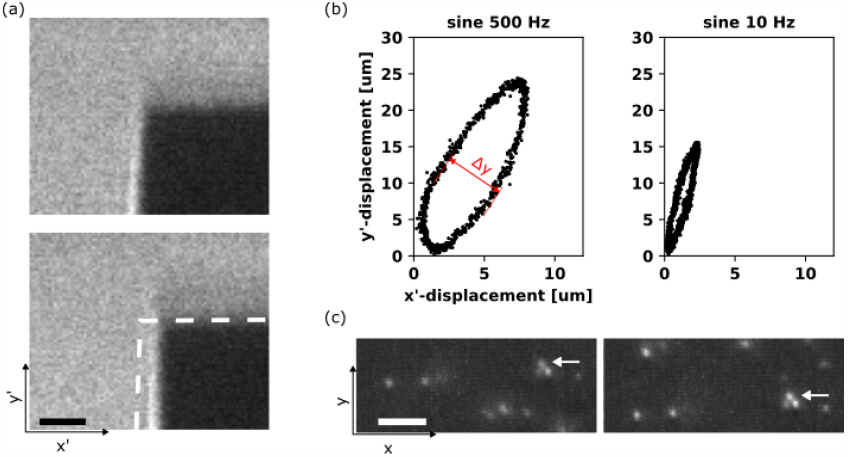
lateral image shift during refocusing: (a) Still frames of a close up high speed video of the edge of the remote retroreflector during high-speed refocusing. The first frame shows the position during the up-stroke, the second frame the position half a period later during the down-stroke. For the same y’-position, the retroreflector is shifted along x’. (b) Quantification of the lateral shift at two different drive frequencies. Higher frequencies lead to a larger shift in x’. (c) Average fluorescence of fluorescent beads during high-speed refocusing showing the same position in the volume during the up-stroke and the down-stroke. Scale bar = 10 μm. The result of the movement quantified in b is a translation of the image along the y axis. Even though the retroreflector shifts around along both dimensions, only the y-component leads to a translation of the actual image since this the axis perpendicular to which the image is folded in the remote focusing plane.

**Figure S1.**
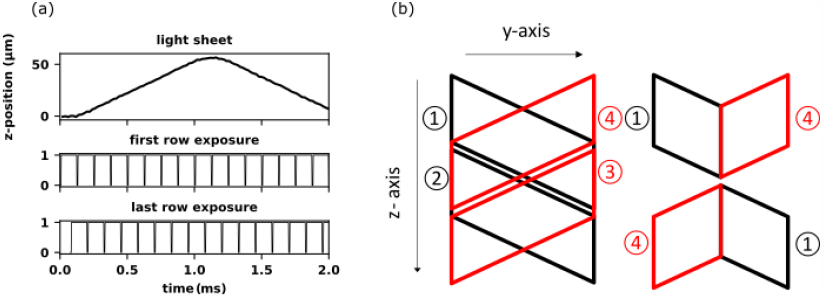
Rolling shutter during fast acquisition: (a) Light sheet position during one period of volume scanning and the exposure timing of the first and last row in the 144 row image. Due to the rolling shutter of the sCMOS camera, the last row starts exposing 76.25 μs after the first row which results in the image being tilted in the recorded volume. (b) Workflow of combining tilted image planes from up- and down-stroke into one consecutive z-stack. The combination of the first half of the first frame of the down-stroke with the second half of the last frame of the up-stroke is combined to become the first image in the z-stack. The second image in the z-stack is combined from the first half of the last frame on the up-stroke and the second half of the first frame of the down-stroke.

**Figure S3.**
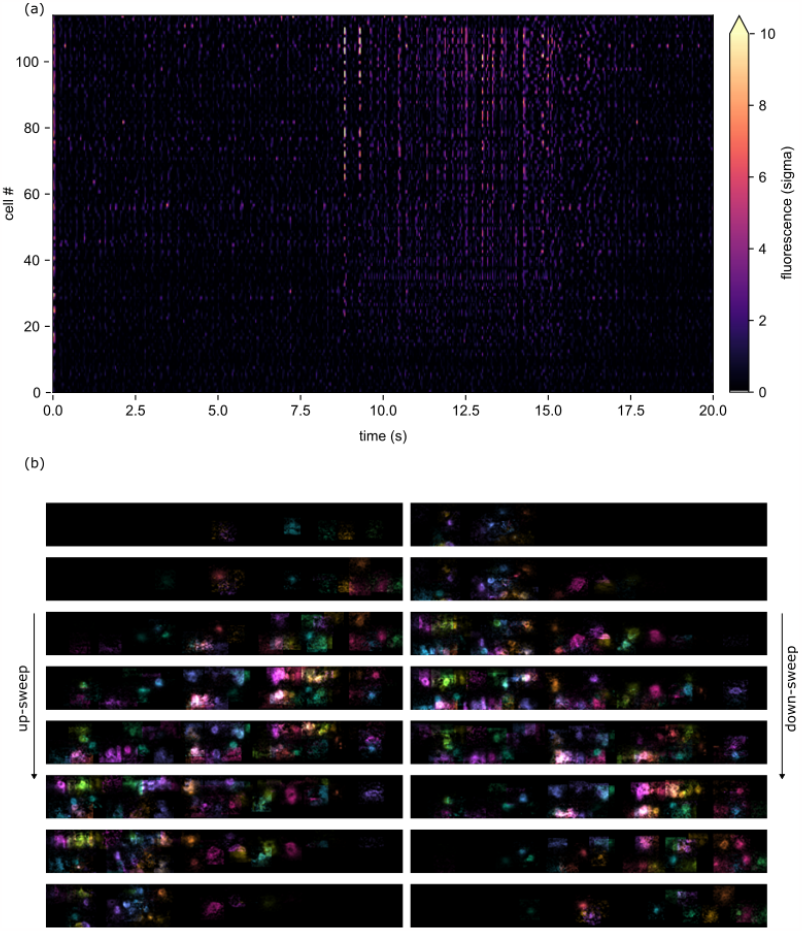
Parallel recording of >100 neurons in a spinal cord volume: (a) Activity of 114 neurons in the recorded volume over 20s. Traces were manually inspected to exclude those that did not show obvious neural activity and close by ROIs that were obviously overlapping. (b) Spatial footprints of the trances shown in a. All 16 frames of one volume sweep are shown with most footprints covering at least two consecutive frames during the up- and the down-sweep.

Video S1 | Sample video of volumetric voltage imaging data. 1s volumetric data (8 planes) spanning the entire volume of the spinal cord. Same dataset as displayed in Fig. 3 and Fig. S3. Gray: Average fluorescent signal over the entire recording. Overlay: Spatial footprints modulated in time by the z-scored ΔF/F of all 114 selected neurons. Overlay adjusted for contrast.

## Notes

### Competing Interest Statement

The authors have declared no competing interest.

